# Cross-Species Evidence for Hippocampal CACNA1C as a Therapeutic Target for Alcohol Use Disorder

**DOI:** 10.1101/2025.09.05.674525

**Authors:** Tanya Pareek, Loc M. Pham, Sinead M. O’Donovan, C. Austin Zamarripa, Obie Allen, Kevin B. Freeman, Donna M. Platt, Kathleen A. Grant, Harry Pantazopoulos, Barbara Gisabella

**Author notes:** **Address all correspondence to:** Barbara Gisabella, PhD, Assistant Professor, Department of Psychiatry and Human Behavior, University of Mississippi Medical Center, 2500 North State Street, TRC 408, Jackson, MS 39216, USA.

## Abstract

Context-induced relapse is a major barrier to recovery from alcohol use disorder (AUD). Identifying molecular targets involved in contextual memories associated with alcohol use may serve as novel pharmacotherapies. Our RNAseq profiling study of the hippocampus from rhesus monkeys with chronic alcohol use identified the voltage-gated calcium channel CACNA1C as a promising therapeutic target. However, data regarding CACNA1C expression in AUD and whether inhibition of CACNA1C can attenuate ethanol contextual memories remains limited. We tested the hypothesis that hippocampal CACNA1C expression is increased in human and nonhuman primates (NHPs) with chronic alcohol use. Further, we used a mouse conditioned place preference (CPP) paradigm to test the hypothesis that Nifedipine, a CACNA1C-selective L-type calcium channel antagonist, can attenuate ethanol-induced CPP. CACNA1C mRNA expression was increased in the hippocampus of subjects with AUD (p<0.03). Increased densities of CACNA1C neurons (p<0.01) and glia (p<0.02) were observed in rhesus monkeys with chronic alcohol use. Ethanol-treated mice spent more time in the ethanol-paired chamber compared to the vehicle animals (p<0.04), demonstrating ethanol-induced CPP. This effect was attenuated by Nifedipine, as time spent in the ethanol-paired chamber in the ethanol + Nifedipine group was not significantly different from the vehicle group. These findings demonstrate that chronic alcohol use increases CACNA1C expression in the hippocampus across species and that a CACNA1C subtype-selective antagonist reduces ethanol-induced CPP. Together, these results support CACNA1C as a promising therapeutic target for context-induced relapse in AUD.

## 1. INTRODUCTION

Alcohol use disorder (AUD) remains a significant public health concern, affecting approximately 28.9 million individuals in the United States ^1^. Despite the availability of efficacious pharmacotherapies for AUD, high relapse rates persist, limiting long-term recovery ^2, 3^. One major factor contributing to relapse is context-induced craving, where environmental cues associated with prior alcohol use trigger compulsive drinking behaviors ^4^. While extensive research has focused on dopaminergic and corticostriatal pathways in addiction, there is increasing recognition that the hippocampus plays a key role in encoding alcohol-associated contextual memories that drive relapse ^5^. The hippocampus is involved in integrating contextual information with reward-related learning, making it a crucial structure in alcohol-associated memory formation ^6^. For example, studies have demonstrated that alcohol exposure enhances hippocampus-dependent learning processes, reinforcing alcohol-associated cues and increasing relapse susceptibility ^7, 8^. However, chronic alcohol exposure has also been shown to disrupt synaptic plasticity and memory function, further complicating extinction learning and relapse prevention ^9^. These findings suggest that alcohol alters hippocampal plasticity in a complex manner (e.g., strengthening maladaptive reward memory processes while impairing general cognitive function). However, the molecular mechanisms underlying these hippocampal adaptations remain unclear. Hippocampal molecular pathways altered with chronic alcohol use may serve as potential therapeutic targets.

L-type voltage-gated calcium channels (LTCC), particularly those encoded by the CACNA1C gene (Cav1.2), may represent molecular therapeutic targets. LTCCs are critical regulators of synaptic plasticity and learning processes within the hippocampus ^10^. Cav1.2-mediated calcium influx is essential for activity-dependent gene transcription and long-term potentiation (LTP), both of which may contribute to alcohol-seeking behavior and relapse ^6, 11^. Furthermore, altered CACNA1C expression has been implicated in psychiatric disorders including major depressive disorder^12–14^, which shared a high degree of co-morbidity with AUD ^15, 16^. Recent evidence also suggests a direct link between CACNA1C, and substance use disorders. For example, pharmacological inhibition of LTCCs in rodents has been shown to reduce alcohol consumption, supporting the idea that Cav1.2 upregulation may contribute to alcohol-seeking behavior ^6^. Additionally, genetic variations in CACNA1C have been associated with altered reward circuitry, which may influence an individual’s susceptibility to addiction ^17^. Despite these findings, there is a lack of information regarding expression of CACNA1C in the primate hippocampus following chronic alcohol use, limiting the development of pharmacological strategies.

Recently, our group identified CACNA1C as a top therapeutic target for reversing transcriptional alterations in the hippocampus of rhesus monkeys following chronic ethanol exposure ^18^, suggesting that increased Cav1.2 expression may contribute to memory impairment and context-induced relapse associated with chronic alcohol use ^19–22^. However, whether these mRNA changes correspond to increased Cav1.2 protein expression and if these changes are region- and/or cell-type-specific remain unknown.

The lack of information regarding CACNA1C expression in the hippocampus of subjects with AUD limits the translation of preclinical findings to clinical populations. Human postmortem studies on subjects with AUD are complicated by inherent confounding factors, such as psychiatric comorbidities and medication exposure. Furthermore, subjects with AUD have a high degree of comorbidity with major depressive disorder (MDD) ^23^. These features are challenging to capture in preclinical models, potentially limiting translatability. We used a cohort of human postmortem samples from subjects with AUD with or without comorbid MDD to test the hypothesis that CACNA1C expression is increased in the hippocampus of subjects with AUD. This cohort includes retrospective clinical assessments, toxicology reports, history of substance use and medication history, allowing for testing of several of the features inherent in the population with AUD. Subjects with comorbid AUD and MDD, and subjects with MDD without AUD were included to examine the potential effects of comorbid MDD on CACNA1C expression in AUD. Furthermore, we used hippocampal tissue collection from nonhuman primates to test the hypothesis that chronic ethanol use increases hippocampal CACNA1C expression in subregions and specific cell types of the hippocampus without the complexity of the confounding factors present in human studies. Lastly, we used a mouse condition place preference paradigm to evaluate whether Nifedipine, a Cav1.2-selective L-type calcium channel antagonist, can attenuate ethanol-induced CPP as first step in testing the potential for L-type calcium channel blockers as therapeutic compounds for mitigating context-induced relapse in AUD.

## 2. MATERIALS AND METHODS

### 2.1 Subjects and Tissue Collection

#### 2.1.1. Human subjects

Hippocampal samples from individuals with a history of either AUD (n=20) or healthy controls (n=20) were obtained from the University of Mississippi Human Postmortem Brain Core (**Table 1, Supplementary Tables S1-16**). All samples in this cohort are tissue blocks flash-frozen in liquid nitrogen vapor containing the hippocampus. All procedures were approved by the Institutional Review Boards of the University of Mississippi Medical Center, Jackson, MS and the University Hospitals Cleveland Medical Center, Cleveland, OH, and are in accordance with the Declaration of Helsinki. Informed consent from the legally defined next-of-kin was obtained for the collection of tissue, medical records and retrospective psychiatric interviews. Structured Clinical Interview for DSM-IV Axis I Disorders was administered by a Master-level social worker to knowledgeable informants of the subjects, as outlined in ^24^. To determine the subjects’ psychopathology, a board-certified clinical psychologist and a board-certified psychiatrist independently reviewed the diagnostic interview scoring notation, the medical examiner’s report, any prior medical records, and a comprehensive narrative that summarized all scores of information about each subject. The social worker, the clinical psychologist, and the psychiatrist reached a consensus on the diagnosis. The cause of death was determined by the medical examiner. Subjects in either group who met DSM-IV criteria for alcohol use disorder (AUD) and/or major depressive disorder (MDD) diagnosis were included for this study. The presence of psychotropic medications and substances of abuse in blood and urine was determined by the medical examiner’s office. Detailed subject demographic information is provided in **Supplemental Tables 1-16**. Lifetime history of substance use was determined from medical records and family interviews and characterized as yes or no based on the history of chronic use for each substance, including alcohol, cocaine, opioids and marijuana. Furthermore, subjective ratings of lifetime alcohol and nicotine intake were determined from medical records and family interviews and rated from 0 (none) to 4 (high). The presence or absence of recent history of calcium channel blockers was determined from medical records and family interviews.

**Table 1.** Basic cohort demographic information for both human and rhesus macaque subjects. Abbreviations: AUD, alcohol use disorder; MDD, major depressive disorder; PMI, postmortem interval, BAC, blood alcohol concentration, 12-month average. Values represent mean +/- standard error.

Human subjects
| Characteristics | AUD (n=20) | MDD (n=20) | MDD/AUD (n=20) | Control (n=20) |
| --- | --- | --- | --- | --- |
| Age | 34.8 (9.0) | 47 (11.4) | 43.9 (11.1) | 43.2 (11.4) |
| Sex | 17M/ 3F | 14M/6F | 16M/8F | 13M/7F |
| pH | 6.5 (0.2) | 6.5 (0.3) | 6.5 (0.3) | 6.5 (0.3) |
| PMI | 18.6 (8.5) | 23.7 (5.9) | 20.2 (7.6) | 21.2 (6.8) |
| Onset of AUD (y/o) | 16.7 (3.3) | N/A | 24.9 (11.9) | N/A |
| Duration of AUD | 20.6 (8.9) | N/A | 20.7 (9.4) | N/A |

Nonhuman Primate Subjects (MATRR Cohort 5)
| Sex | Age | Group | Drinking Days | Drinking Category |
| --- | --- | --- | --- | --- |
| M | 7 | EtOH | 364 | BD |
| M | 7 | EtOH | 364 | HD |
| M | 7 | EtOH | 365 | HD |
| M | 7 | EtOH | 356 | HD |
| M | 7 | EtOH | 365 | VHD |
| M | 7 | EtOH | 364 | VHD |
| M | 7 | EtOH | 363 | VHD |
| M | 7 | EtOH | 361 | VHD |
| M | 8 | Control | 0 | N/A |
| M | 8 | Control | 0 | N/A |
| M | 6 | Control | 0 | N/A |
| M | 5 | Control | 0 | N/A |
| M | 5 | Control | 0 | N/A |

Binge Drinker (BD), Heavy Drinker (HD), Very Heavy Drinker (VHD), Ethanol (EtOH)

Information regarding potential confounding variables (see **Supplemental Tables 1-16**) including time of death calculated as zeitgeber time (ZT), postmortem interval (PMI), tissue pH, race, age and sex, history of substance use, pharmacological treatment, sleep quality, suicide, duration of AUD, duration of major depression, and mood symptom severity in the last two weeks of life were obtained from medical records, police reports, medical examiner’s reports, and family interviews. Toxicology reports were used to determine the presence or absence of alcohol, cocaine, opioids, SSRIs, antidepressants, antipsychotics, lithium and benzodiazepines in the blood at death. Sleep quality was determined by subjective rating of chronic history of sleep disturbances or history of excessive sleep reported in medical records and family interviews. Mood symptom severity in the last two weeks of life was determined from medical records and family interviews. Sample locations were identified using the Atlas of the Human Brain ^25^. All procedures were approved by the Institutional Review Boards of the University of Mississippi Medical Center and the University Hospitals Cleveland Medical Center, in accordance with the Declaration of Helsinki.

#### 2.1.2. Rhesus monkeys

Fresh, frozen hippocampal samples from adult, male rhesus monkeys (*Macaca mulatta*), with a history of chronic, oral ethanol use (n=7) or no alcohol (n=5) were used (**Table 1**). Briefly, monkeys were experimentally naïve at the onset of alcohol induction and followed an open-access protocol ^26^. Open-access (i.e., 22 hours/day) self-administration sessions occurred daily for 12 months, with average ethanol intake ranging between 2.5 and 3.8 g/kg/day. Cohort 5 monkeys used were grouped as heavy drinkers (HD; n=3) or very heavy drinker (VHD; n=4). Control monkeys’ self-administration sessions were identical to those orally self-administering alcohol. However, the control monkeys had access to water and 10% maltose dextrin solution. Samples were obtained from the Monkey Alcohol Tissue Research Resource (MATRR) Cohort 5 (www.matrr.com). Here, following the 12-month open access period and no abstinence (i.e., within 4-8 hrs of last self-administration session end), tissue was collected ^27^. All animal-use procedures were approved by the Oregon National Primate Research Center’s Institutional Animal Care and Use Committee and were conducted in accordance with the National Research Council’s Guide for Care and Use of Laboratory Animals (8th edition, 2011).

#### 2.1.3

Adult, male mice (C57BL/6J; The Jackson Laboratory) (n=10/group) began studies at approximately 70-80 days of age. To acclimate and reduce stress, mice were pair-housed in cages for three weeks before the start of the studies. Animals were maintained in a vivarium under a 12-hour reverse light/dark cycle with lights off at 0700 hours and provided with ad libitum food and water. All studies were conducted during the dark cycle under infrared lighting conditions. All animal-use procedures were approved by the University of Mississippi Medical Center’s Animal Care and Use Committee and were conducted in accordance with the National Research Council’s Guide for Care and Use of Laboratory Animals (8th edition, 2011).

### 2.2 Quantitative real-time polymerase chain reaction (qRT-PCR)

The gene expression of CACNA1C mRNA was measured from human postmortem hippocampal samples using qRT-PCR. RNA was extracted from 14 µm hippocampal cryosections using the RNeasy Minikit (cat# 74104, Qiagen, NL) according to the manufacturer’s instructions. On-column DNAse digestion was carried out using the RNAse-free DNAse set (#79254, Qiagen, NL), per manufacturer’s guide. Complementary DNA (cDNA) was synthesized using a High-Capacity cDNA Reverse Transcription Kit (Applied Biosystems, Foster City, CA, USA). For each reaction, 0.5 μl of cDNA (1:3 diluted) was placed in a 10 μL reaction containing 5 μL of SYBR Green PCR Master Mix (Applied Biosystems) and 10 pmol of each primer (Invitrogen, United States). The primers used are listed in **Supplemental Table 17.**

Custom-made assays were tested for specificity and resulted in a single band of expected size. All reactions were performed in triplicate using 384-well optical reaction plates (Life Technologies, United States) on an Applied Biosystems detection system (QuantStudio 5, Applied Biosystems, Life Technologies, United States). Reactions were performed with an initial ramp time of 10 min at 95°C, and 40 subsequent cycles of 15 s at 95°C and 1 min at 60°C. For negative controls for the qPCR reactions, non-template control (cDNA was omitted) and no-RT control (reverse transcriptase excluded from cDNA synthesis reaction) were run on each plate. Relative concentrations of the transcripts of interest were calculated with a comparison to a standard curve made with dilutions of cDNA from a pooled sampling of all the subjects. Values for the transcripts of interest were normalized to the geometric mean of B2M, ACTB, GAPDH and PPIA values for the same samples. Data were collected by QuantStudio Design and Analysis software v1.5.1.

### 2.3 Immunohistochemistry and Immunofluorescence

CACNA1C Immunoreactive neurons and glia were evaluated using immunohistochemistry and immunofluorescence. Hippocampal samples from monkeys that were fresh, frozen and cryosectioned at 30 micrometers on glass slides were used.

#### 2.3.1 Immunohistochemistry for CACNA1C

Fresh-frozen, slide-mounted medial hippocampal cryosections from rhesus macaque subjects (n= 3-5 sections/animal) were post-fixed for 30 minutes in 4% paraformaldehyde, followed by 1-hour incubation in 2% bovine serum albumin (BSA), and overnight incubation in rabbit anti-CACNA1C (ProteinTech, cat# 21774-1-AP, 1:500). The following day, tissue was incubated in streptavidin horseradish peroxidase (1:3000 μl, Zymed, San Francisco, CA) for four hours, followed by a 20-minute incubation in 3’3-diaminobenzidine with nickel sulfate hexahydrate and H2O2. The tissue was dehydrated through an ethanol and xylene series, followed by histological labeling with Methyl Green (catalog #H-3402, Vector Labs). Sections were coverslipped and coded for blinded quantitative analysis. All sections included in the study were processed simultaneously to avoid procedural differences.

#### 2.3.2 Immunofluorescence for CACNA1C and Excitatory Neurons (CACNA1C and CamKIIα Labeling)

Slides (n=3/animal) were post-fixed with 1 mL PFA (4°C) per slide for 30 minutes, followed by a PB wash (5 minutes) and three washes in PBS-Tx (5 minutes each). Endogenous peroxidase was blocked with 0.3% H₂O₂/10% methanol in PBS-Tx for 30 minutes, followed by a 1-hour incubation in 2% BSA. Primary antibody incubation was performed overnight at room temperature in 2% BSA containing rabbit anti-CACNA1C (ProteinTech, cat# 21774-1-AP, 1:500) and goat anti-CamKIIα (Invitrogen, cat# PA5-19128, 1:300) or mouse anti-GFAP (BioLegend, cat# 837201, 1:500). The following day, slides were washed and incubated for 4 hours in goat anti-rabbit Alexa Fluor 555 (1:300) and goat anti-mouse Alexa Fluor 647 (1:300) for CamKIIa or goat anti-mouse 647 (1:300) for GFAP in 2% BSA, followed by incubation in DAPI (1:5000 in PB) for 20 minutes and incubation in Trueblack lipofuscin autofluorescence quencher (cat#23007, Biotum) for 2 minutes. Sections were mounted on gelatin coated glass slides and coverslipped with Dako fluorescent mounting media (cat#S3023, Agilent Technologies).

### 2.4 Imaging and Quantification

Quantification was performed using an Olympus BX61 microscope and CACNA1C-labeled neurons and glia were distinguished based on morphology. Each anatomical subdivision of the hippocampus was traced for an area measurement using a 4X objective. Each subregion was systematically scanned through the full x, y, and z-axes under 40x magnification using Stereo-Investigator v11 to quantify all CACNA1C-labeled cells within each subregion.

### 2.5 Apparatus and CPP Procedural sequence

Five sound-attenuated 3-compartment chambers equipped with manual doors, lights, black and white guillotine doors, and stainless-steel floor (Med Associates Inc, St. Albans, VT, USA) were used for all sessions. The chambers were equipped with infrared photobeam strips in each compartment to measure locomotor activity and location (i.e., location, activity count and time spent in each chamber). A PC equipped with Med-Associates software (Med-PC for Windows; St. Albans, VT, USA) controlled experimental conditions and recorded data.

Animals were handled and habituated to injections for one week before the start of the 12-day CPP protocol. The CPP design consisted of a pre-test assessment, eight days of drug-paired conditioning and one post-test assessment. On Days 1-3, mice underwent a 20-minute pre-test session in which time spent in each chamber was recorded to determine initial preference. For drug assignments, a biased CPP design was used, meaning each animal was assigned to receive the experimental condition in its non-preferred chamber ^28^. For treatment assignments, groups were determined based on the 3-day average time spent in the black chamber. The non-preferred chamber was then assigned to one of four treatment groups: saline + vehicle, ethanol + vehicle, saline + Nifedipine, or ethanol + Nifedipine. All conditions were counterbalanced. During the 8-day conditioning phase, animals underwent a 5-minute conditioning session each day where they received either vehicle or drug injections. On Days 4, 6, 8 and 10, animals received two vehicle injections and were confined to the preferred chamber. On Days 5, 7, 9 and 11, animals received their assigned experimental conditions described above, and animals were confined to the non-preferred chamber. For all conditioning days, all pretreatments were administered intraperitoneally (i.p.). Nifedipine (5 mg/kg; i.p.) and Nifedipine vehicle (2% DMSO + 1% Tween80) were administered fifteen minutes prior to the start of the conditioning session. Ethanol (2 g/kg, 26.5% v/v; i.p.) and saline pretreatments were administered two minutes prior to the conditioning session. The dose of nifedipine and ethanol were based on previous studies showing them to be suitable for CPP studies in mice (^29–32^. On Day 12, animals underwent a 20-minute post-test assessment (i.e., identical to pre-test conditions) to reestablish preference scores.

### 2.6 Drugs and pretreatment

For the pretreatments, ethanol (95%; Pharmco Products, Brookfield, CT, USA) was diluted to 2 g/kg, 26.5% *v/v*). The solution was prepared using 0.9% saline. Nifedipine and Dimethyl sulfoxide (DMSO) were purchased from Sigma-Aldrich (St. Louis, MO, USA). Nifedipine was prepared in 100% Dimethyl sulfoxide (DMSO) and diluted to 2% DMSO using 1%Tween80 and saline. 2% DMSO + 1% Tween 80 was used as the Nifedipine vehicle. The Nifedipine solution was prepared daily for drug conditioning sessions. Ethanol and Nifedipine vehicle solutions were used for up to 30 days before being replaced with new solution. All solutions were filtered with 200 µm filters before being administered intraperitoneally. All pretreatments were administered i.p. at a volume of 0.24-0.38 mL.

### 2.7 Data Analysis

#### 2.7.1. Human postmortem mRNA analysis

qRT-PCR mRNA values were obtained using QuantStudio Design and Analysis Software v1.5.1. Differences between groups relative to the main outcome measures were assessed for statistical significance using stepwise linear regression analysis of covariance (ANCOVA) using JMP Pro v16.1.0 (SAS Institute Inc., Cary, NC). Logarithmic transformation was applied to values when not normally distributed. Potential confounding variables (**Table 1 and Supplementary Tables 1-16**) were tested systematically for their effects on main outcome measures and included in the model if they significantly improved goodness-of-fit. Covariates found to significantly affect outcome measures are reported. Subjects with AUD, subjects with AUD/MDD and subjects with MDD only were compared separately with psychiatrically normal controls.

#### 2.7.2. Rhesus monkey microscopy data analysis

Numerical densities of microscopy measures were calculated as described in our previous studies ^33–35^. Differences between groups relative to the main outcome measures were assessed for statistical significance using stepwise linear regression analysis of covariance (ANCOVA) using JMP Pro v16.1.0 (SAS Institute Inc., Cary, NC). Logarithmic transformation was applied to values when not normally distributed.

#### 2.7.3. Mouse Behavioral Analysis

Each cohort was designated according to the between-subject condition of Drug Treatment (saline + vehicle, ethanol + vehicle, saline + Nifedipine, ethanol + Nifedipine). Primary outcomes were changes in preference score, activity, and movement. Change-from-baseline values were expressed using the following formula:

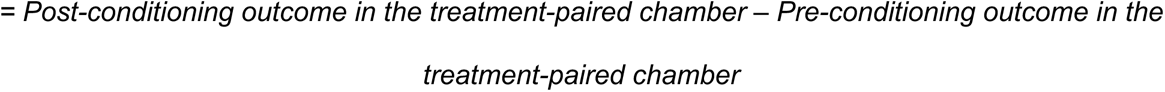

where the treatment was either the drug (i.e., ethanol, Nifedipine, or ethanol and Nifedipine) or vehicle. Within each group, change in time spent, activity and movement in the treatment-paired chamber were average across mice, and data were analyzed using three one-way analysis of variances (ANOVA) with the between-subjects factor of Drug Treatment (saline + vehicle, ethanol + vehicle, saline + Nifedipine, ethanol + Nifedipine). Planned comparisons (Fischer’s LSD tests) were used to compare each condition to the vehicle control group. For all analyses, statistical significance was defined as an alpha level of < 0.05. Analyses were conducted using GraphPad Prism, Version 9 (La Jolla, CA).

## 3. RESULTS

### 3.1 Increased expression of CACNA1C in the hippocampus of subjects with Alcohol Use Disorder

Significantly increased CACNA1C mRNA expression was observed in the hippocampus of subjects with AUD, adjusted for significant effects of exposure to calcium channel blockers, ZT time of death, and PMI (**Figure 1**). Similar increased expression was detected in subjects with comorbid AUD/MDD (p<0.05) adjusted for significant effects of exposure to calcium channel blockers and tissue pH. In comparison, subjects with MDD only displayed significantly lower CACNA1C mRNA expression compared to control subjects (p<0.04), adjusted for significant effects of exposure to calcium channel blockers, ZT time of death, PMI and presence of SSRIs in the blood at death (**Figure 1**).

**Figure 1.**
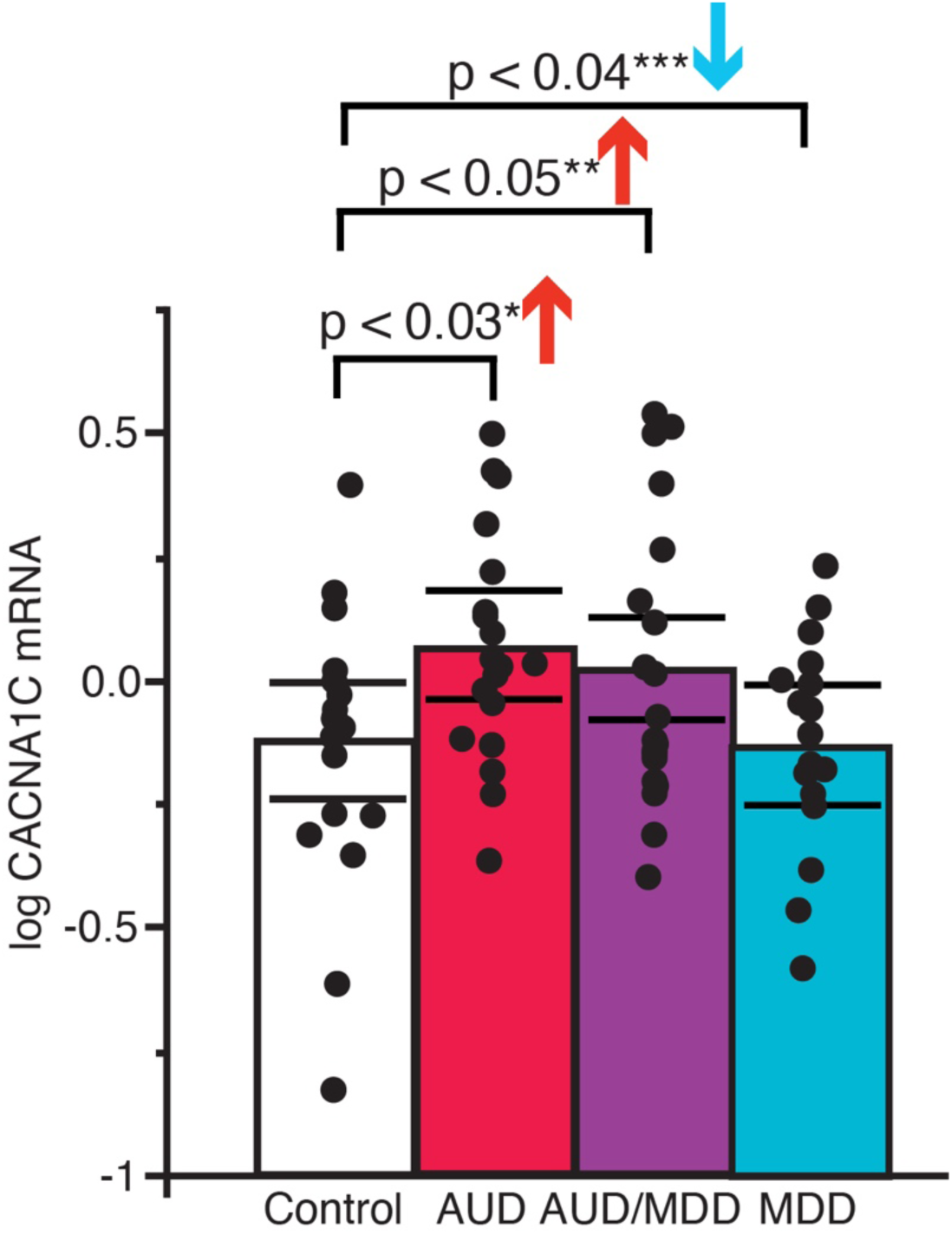
Increased CACNA1C mRNA expression in the hippocampus of subjects with AUD. Increased CACNA1C mRNA expression in subjects with AUD, AUD+ MDD and MDD only compared to control subjects (p<0.03, *adjusted for significant effects of exposure to calcium channel blockers, ZT time and PMI). ** adjusted for significant effects of exposure to calcium channel blockers and tissue pH. *** adjusted for significant effects of exposure to calcium channel blockers, ZT time of death, PMI and presence of SSRIs in the blood at death. Bar graph represents mean/group; black lines represent 95% confidence intervals (Cis) and black dots represent individual value for each individual.

### 3.2 Increased densities of CACNA1C cells in the hippocampus of monkeys with chronic alcohol use

#### 3.2.1. Neurons

CACNA1C immunolabeling was observed in neurons across all hippocampal sectors of both monkeys with no alcohol use (**Figure 2A-C)**). Significantly greater densities of CACNA1C neurons were observed in the hippocampus of monkeys with chronic ethanol use compared to control subjects (*p*<0.01); **Figure 2D**). Greater densities of CACNA1C neurons were present across hippocampal sectors, displaying statistical significance in the majority of hippocampal sectors (i.e., CA3 stratum oriens (*p*<0.05); CA3 stratum pyramidale (*p*<0.005); CA2 stratum pyramidale (*p*<0.001); CA1 stratum pyramidale (*p*<0.03)) of the ethanol group when compared to the control group (**Figure 2E**).

**Figure 2.**
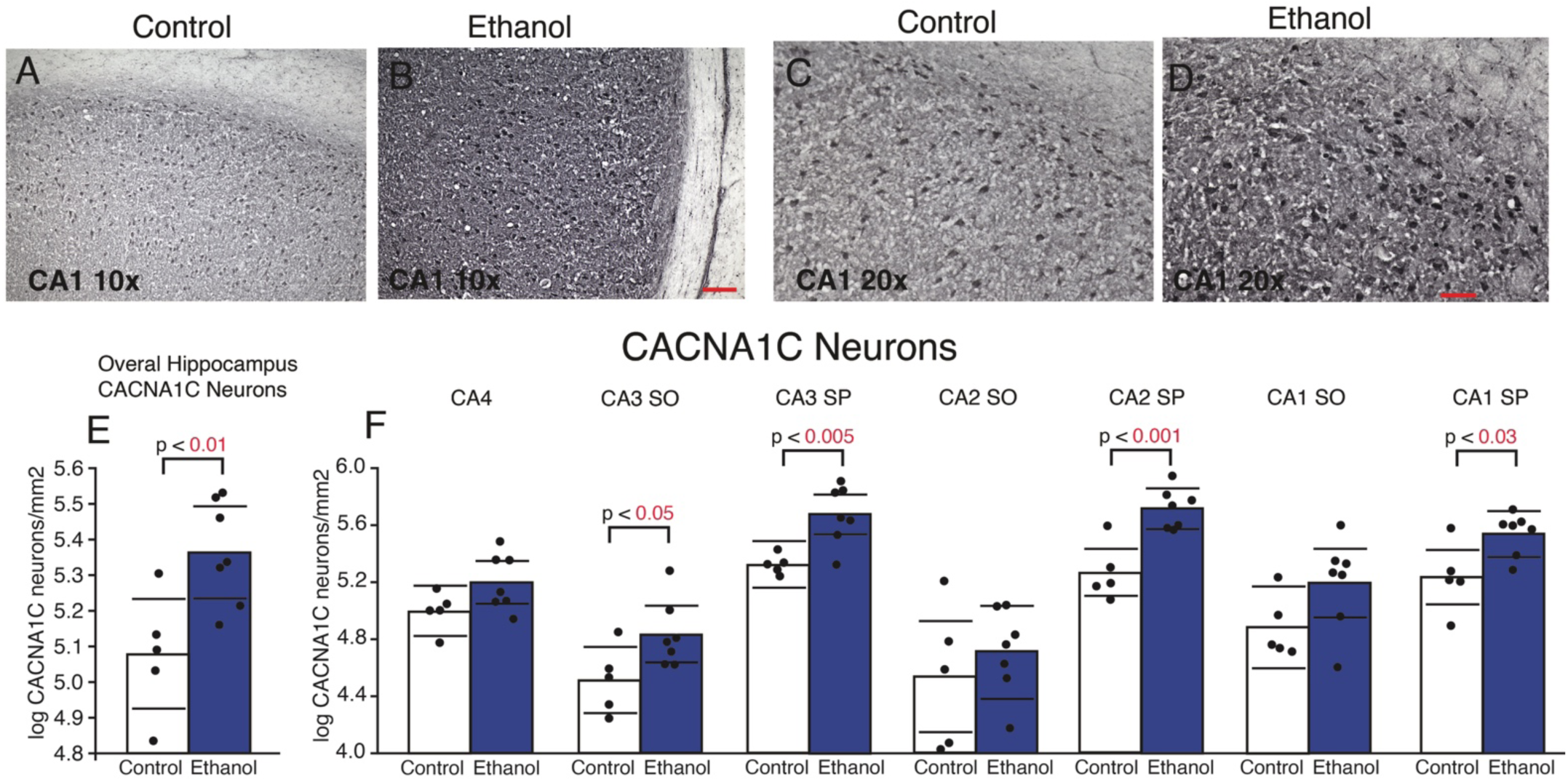
Increased Densities of CACNA1C Neurons in the hippocampus of monkeys with chronic ethanol administration. Representative 10x magnification images of CACNA1C labeling in control animals **(A),** and animals chronic ethanol use **(B)**. 20X images of CACNA1C expression in neurons in CA1 sector of a monkey with no alcohol use **(C)** and chronic use **(D).** Scale bars = 100 μm. Total CACNA1C densities between control and ethanol groups, x-axis represents group and y-axis represents density of CACNA1C neurons/square millimeter of tissue **(D)** and within hippocampal sectors, where x-axis represents groups and hippocampal regions and y-axis represents density of CACNA1C+ neurons/square millimeter of tissue **(E)** Bar graphs represent mean/group; black lines represent 95% confidence intervals (Cis) and black dots represent individual value for each monkey

#### 3.2.2. Glia

We observed CACNA1C immunolabeling in cells with glial morphology across all hippocampal subregions. Densities of CACNA1C glia were significantly increased in monkeys with chronic ethanol use compared to control subjects (*p*<0.02); **Figure 3A**). CACNA1C glial densities were increased in all hippocampal regions, attaining significance in CA3 stratum oriens (p<0.05), CA3 stratum pyramidale (p<0.03), CA2 stratum oriens (p<0.004), CA2 stratum pyramidale (p<0.006), and CA1 stratum pyramidale (p<0.005) (**Figure 3B**).

**Figure 3.**
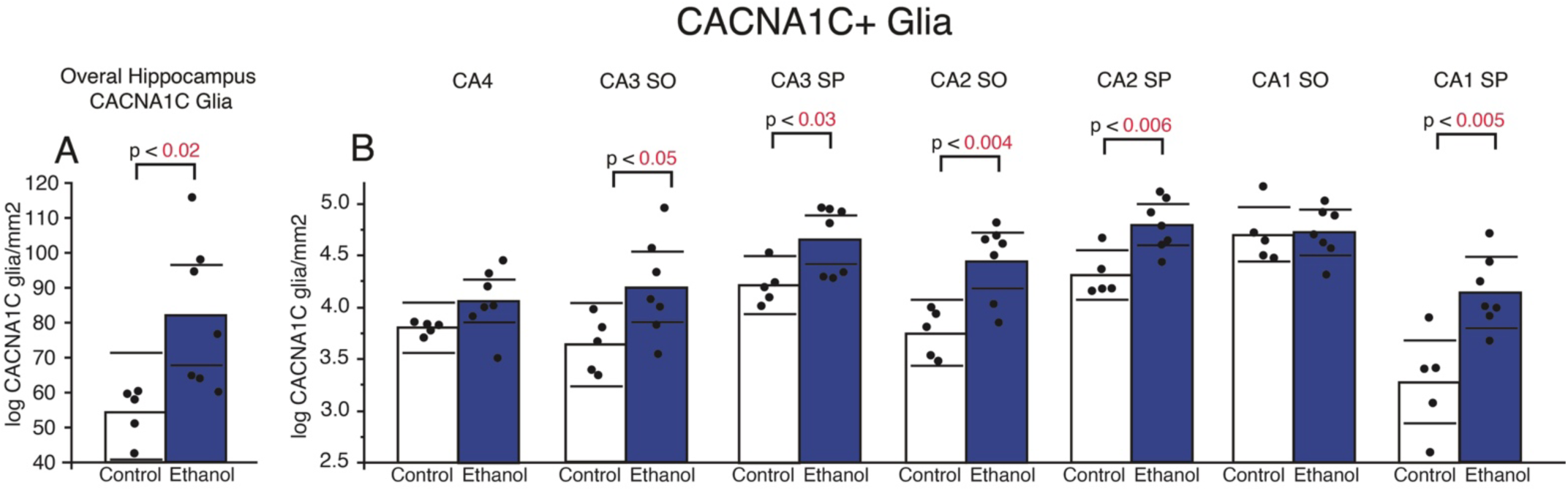
Increased Densities of CACNA1C Glial Cells in the hippocampus of monkeys with chronic ethanol administration. Total CACNA1C densities between control and ethanol groups x-axis represents groups and y-axis represents density of CACNA1C glia/square millimeter of tissue **(A)** and within hippocampal sectors where x-axis represents groups and hippocampal regions and y-axis represents density of CACNA1C glia/square millimeter of tissue **(B).** Bar graphs represent mean/group; black lines represent 95% confidence intervals (Cis) and black dots represent individual value for each monkey

### 3.3 CACNA1C expression corresponds to excitatory neurons and astrocytes

We used co-labeling of CACNA1C with the excitatory neuron marker CamKIIα to test the hypothesis that CACNA1C immunoreactive neurons correspond primarily to excitatory neurons (**Figure 4**). CACNA1C positive neurons primarily colocalized with CamKIIα neurons (i.e., 62%) (**Figure 4D**). We used co-labeling of CACNA1C with GFAP (**Figure 4**), to test the hypothesis that CACNA1C glial cells correspond to astrocytes. CACNA1C positive glial cells partially co-localized with GFAP positive astrocytes (i.e., 36%) (**Figure 4H**).

**Figure 4.**
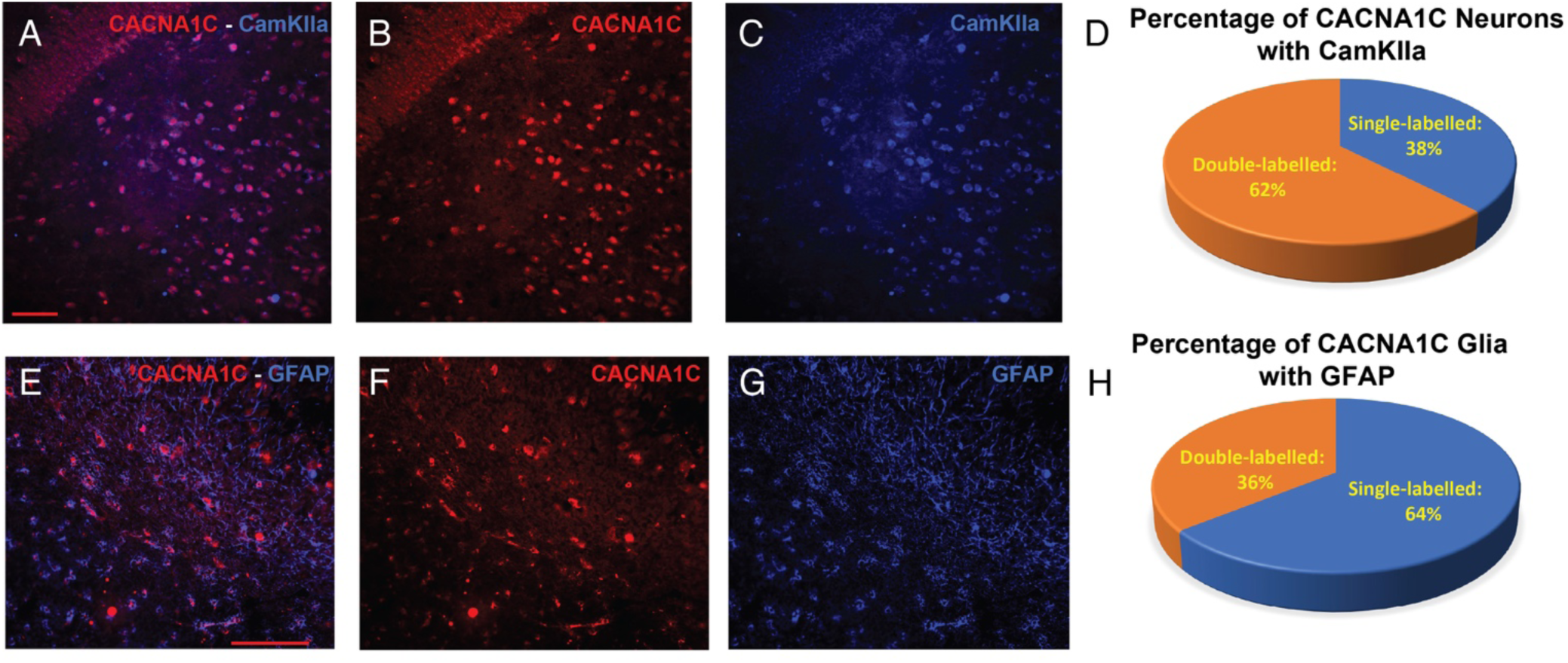
CACNA1C Expression in Excitatory Neurons and Astrocytes. CACNA1C and CamKIIa co-labeled neurons **(A)** CACNA1C labeled neurons **(B)** and CamKIIa labeled excitatory neurons **(C).** Overall percentage of CACNA1C single and double labeling with CamKIIa **(D);** CACNA1C and GFAP co-labeled glia **(E)**, CACNA1C labeled glia **(F)**, and GFAP labeled astrocytes **(G).** Overall percentage of CACNA1C single and double labeling with GFAP **(H).**

### 3.4 Attenuation of ethanol-induced CPP in male mice by Nifedipine

Experimental timeline of CPP studies is depicted in **Figure 5A** Change-from-baseline values are illustrated in **Figure 5** for Time (**5B**), Activity (**5C**), and Movement (**5D**) in the drug-paired chamber. There was no main effect of Drug Treatment observed for the ANOVA for time spent in the drug-paired chamber. Planned comparisons between the vehicle group and the experimental conditions revealed that the ethanol + vehicle group (**Figure 5B**) spent significantly more time in the ethanol-paired chamber relative to vehicle group (*p* = 0.0458). In contrast, these comparisons revealed that both the Nifedipine and ethanol + Nifedipine groups exhibited no changes in time spent in the drug-paired chamber relative to the vehicle group (**Figure 5B**). There was no significant main effect of Drug Treatment for both ANOVA results for both activity and movement (**Figure 5C** and **2D**). Further, there were no significant differences observed in activity and movement for any of the planned comparisons relative to vehicle alone (**Figure 5C** and **5D**).

**Figure 5.**
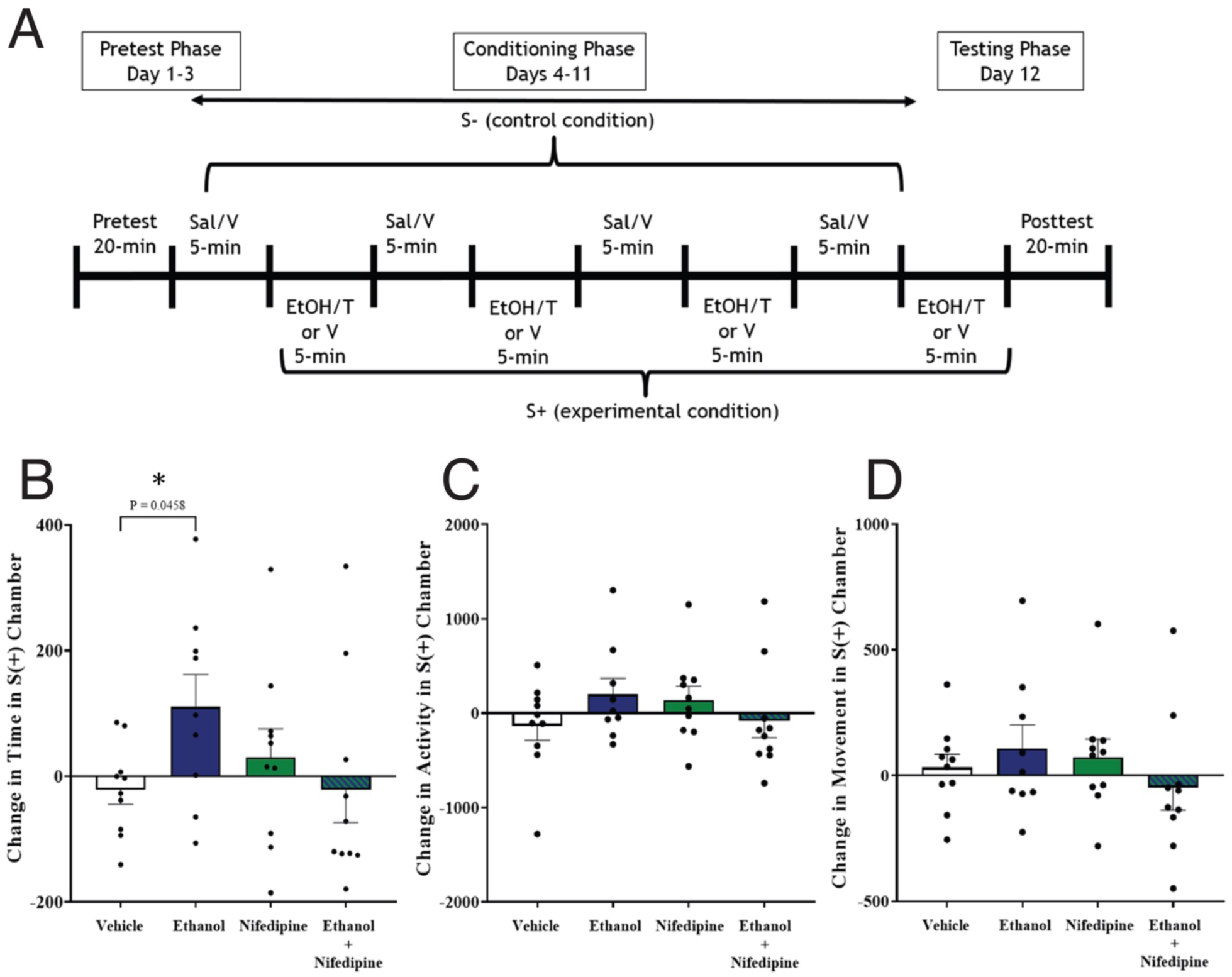
The L-type calcium channel antagonist nifedipine blocks ethanol induced conditioned place preference. **(A)** Overall procedural sequence consisting of a pre-test phase (Day 1-3); a conditioning phase (Days 4-12) where S-represents the control condition (i.e., Saline/ Nifedipine Vehicle) and S+ represents the experimental condition (i.e., Nifedipine/Nifedipine Vehicle + Ethanol/Saline); a post-test phase (Day 12. Times represent the length of the session per day. X-axis represents the treatments groups and y-axis represents change in time spent in the S+ chamber **(B),** change in activity in the S+ chamber **(C)** and change in movement **(D).** *Denotes a significant difference from vehicle group

## 4. DISCUSSION

Our findings represent the first evidence for increased CACNA1C mRNA expression in the hippocampus of subjects with AUD and subjects with comorbid AUD+MDD. Furthermore, our findings of decreased CACNA1C expression in subjects with MDD-only in comparison to the increased expression in subjects with AUD and comorbid AUD+MDD (**Figure 1**) highlights the importance of considering comorbidities when examining neuropathology of these disorders and designing novel pharmacological interventions. Increased CACNA1C protein expression in hippocampal neurons and glial cells following chronic ethanol use in rhesus monkeys **(Figures 2 and 3)** provide further support for increased CACNA1C expression in subjects with AUD. CACNA1C neuronal expression predominantly in excitatory neurons (CamKIIα+) (**Figure 4D**) supports previous reports in rodent brain ^6^. In addition, we observed previously unreported CACNA1C expression in glial cells which partially colocalized with the mature astrocyte marker GFAP (**Figure 4H**), suggesting that chronic alcohol use impacts CACNA1C expression in several cell types. Lastly, our mouse behavioral findings suggest that pharmacological inhibition of the CACNA1C subtype can reduce the context-dependent rewarding effects of ethanol, highlighting its potential as a promising therapeutic target for preventing context-induced relapse in AUD. Together, these findings provide the first cross-species evidence that chronic alcohol use increases CACNA1C expression in the hippocampus, a molecular adaptation that may contribute to context-induced relapse vulnerability.

Our observed increase of CACNA1C mRNA expression in the hippocampus of subjects with AUD aligns with previous preclinical studies demonstrating that chronic alcohol exposure upregulates CACNA1C expression in the rodent hippocampus ^6^. Consistent with these findings, CACNA1C’s critical role in synaptic plasticity and learning processes suggests that its upregulation in AUD may reflect a state of heightened hippocampal excitability and maladaptive memory persistence ^36, 37^. Speculatively, this adaptation may facilitate the strengthening of alcohol-associated contextual memories and contribute to relapse vulnerability. Importantly, our findings provide key insight into CACNA1C expression in comorbid AUD+MDD and MDD only. Increased CACNA1C expression in subjects with AUD+MDD compared to reduced expression in subjects with MDD-only suggests that increased CACNA1C expression may be distinct to substance use pathology that may partially counteract decreased expression in MDD. Preclinical evidence that decreased CACNA1C expression during embryonic development increases the susceptibility to stress suggests that decreased CACNA1C expression may stem from genetic factors (e.g., predispose people to developing MDD and/or AUD following stressful experiences ^38^). Reports demonstrating no change in CACNA1C expression in the mouse hippocampus following chronic unpredictable stress suggest that decreased hippocampal expression in subjects with MDD may develop from genetic factors ^39^. The disease-specific differences in CACNA1C expression in our study highlight the importance of considering comorbid conditions when developing novel pharmacological approaches. Moreover, these findings highlight key neurobiological distinctions between AUD and MDD, reinforcing that addiction and depression involve overlapping but distinct mechanisms ^40, 41^. Increased CACNA1C protein expression in rhesus monkeys following chronic ethanol use provides further support for the increased CACNA1C expression in subjects with AUD in the nonhuman primate hippocampus without the concerns of potential confounding factors such as comorbid MDD, exposure to other drugs of use, and exposure to pharmacological therapies inherent with human postmortem studies. Furthermore, our findings represent the first evidence of increased CACNA1C protein expression in a nonhuman primate model of AUD.

The cell-type specificity of CACNA1C upregulation provides further mechanistic insight into its role in AUD-related neuroplasticity. The majority of CACNA1C-expressing neurons were CamKIIα+, indicating selective upregulation in excitatory neurons. This is consistent with CACNA1C’s role in synaptic plasticity and LTP, which are key processes involved in contextual memory encoding ^42^. Additionally, CACNA1C expression was detected in GFAP+ astrocytes, suggesting that alcohol-induced neuroadaptations extend beyond neurons. Astrocytes play a crucial role in calcium signaling, synaptic maintenance, and neuroimmune regulation ^43, 44^. The observed upregulation of CACNA1C in astrocytes suggests that chronic alcohol exposure may contribute to neuroinflammatory processes, altered glutamate homeostasis and hippocampal hyperexcitability. Several studies have reported decreased astrocytes and astrocytic markers, along with atrophic morphological alterations in astrocytes in subjects with AUD ^45^. This includes reduced astrocyte numbers in the hippocampus of subjects with AUD ^46^. Therefore, atrophic changes and decreased astrocyte numbers reported in subjects with AUD may reflect excitotoxic effects in part from chronically elevated CACNA1C expression. Alternatively, taken together with reports that alcohol impairs astrocyte synthesis and maturation ^47, 48^ and low percentage of colocalization of CACNA1C glia with GFAP in our study (**Figure 4H**), increased densities of CACNA1C glia in monkeys with chronic alcohol use may represent expression in immature glial cells stemming from impaired astrocyte maturation. Future studies examining glial CACNA1C signaling in various glial cell populations and its role in addiction-related neuroplasticity may provide insight into these possibilities.

One major implication of these findings is that CACNA1C upregulation in the hippocampus may augment relapse risk by strengthening alcohol-associated memories. Here, the hippocampus plays a central role in contextual memory encoding, and maladaptive plasticity within this region may contribute to persistent cue-induced alcohol craving ^49^. This aligns with studies showing that alcohol-associated environmental cues can trigger compulsive drinking behaviors ^4^. Further, CACNA1C regulates calcium influx into neurons, an essential process for various processes (e.g., synaptic remodeling and communication) ^50, 51^. Studies have shown that chronic alcohol exposure may induce CACNA1C upregulation as a compensatory adaptation, leading to increased hippocampal excitability and reinforcement of alcohol-context associations ^6, 52^. Similarly, an upregulation in CACNA1C expression in astrocytes may contribute to neuroinflammatory responses, further exacerbating synaptic dysfunction and plasticity ^53^.

Given CACNA1C’s role in hippocampal plasticity and alcohol-related neuroadaptations, our findings suggest that L-type calcium channel antagonists (commonly referred to as CCBs) may have therapeutic potential for reducing alcohol-seeking behavior. Specifically, the role of CACNA1C in alcohol-induced neuroadaptations has been illustrated by the modulation of ethanol’s behavioral effects by CCBs ^54^. This is highlighted by our recent RNAseq study which identified calcium channel blockers as a top discordant mechanism of action ^18^. Our evidence that the L-type calcium channel antagonist nifedipine blocks ethanol induced CPP suggests that L-type calcium channel antagonists may serve as efficacious treatments to mitigate the upregulation of CACNA1C in the hippocampus and weaken contextual memory associated with alcohol use. In various studies, LTCC subtype-selective antagonists (e.g., isradipine, nifedipine) have been shown to reduce drug-seeking behaviors ^29, 55^. Mechanistically, the attenuation of ethanol-induced CPP by Nifedipine likely stems from the blockade of Cav1.2 subtype calcium influx, which is critical for synaptic plasticity, long-term potentiation and context-dependent memory formation ^40, 56^. Previous studies using CPP paradigms have similarly demonstrated that Nifedipine reduces drug reward, supporting the broader role of calcium-dependent mechanisms across substances of misuse ^29^. Although Nifedipine exhibits greater selectivity for Cav1.2 channels, some residual activation on Cav1.3 channels may persist ^57, 58^, potentially influencing the overall effect of ethanol and warranting further investigation to better delineate specific channel contributions.

While these findings provide novel insights into CACNA1C’s role in AUD, several limitations should be considered. First, human postmortem studies inherently present challenges, such as potential confounding variables related to medication exposure and psychiatric comorbidities. While our approach of including subjects with and without comorbid MDD, and confirmation with nonhuman primate studies addressed several of these limitations, and our analysis of covariates in human postmortem subjects accounted for CCB use, future studies examining specific effects of potential confounding factors such as exposure to CCBs or antidepressant treatment may provide critical insight into the nature of increased CACNA1C expression AUD and guide pharmacological strategies. Additionally, we observed hippocampal CACNA1C upregulation at both the mRNA and protein levels, but the impact on hippocampal neuronal activity remains unclear. Future studies integrating electrophysiological approaches are needed to determine how CACNA1C upregulation affects hippocampal synaptic plasticity and memory circuits associated with AUD.

In summary, our results identified increased hippocampal CACNA1C expression in subjects with AUD and nonhuman primates with chronic ethanol use, encompassing neuronal and glial cell populations. Furthermore, L-type calcium channel antagonists such as nifedipine demonstrate potential for mitigating context-induced relapse in AUD. Increased CACNA1C expression in both subjects with AUD and comorbid MDD compared to decreased expression in subjects with MDD only, suggests disease-specific effects. Therefore, the distinction serves as a guide for the development of pharmacotherapeutic strategies.

## ACKNOWLEDGMENTS

The authors want to extend their deepest gratitude to the families consenting to donate brain tissue, the staff of the Cuyahoga County Medical Examiner’s Office, Cleveland, Ohio, the expert assistance (i.e., in the establishment of psychiatric diagnoses, acquiring written consent and/or the collection of the tissues) of Dr. James C. Overholser, Dr. George Jurjus, Dr. Lisa C. Konick, Lesa Dieter, Timothy M. De Jong, Lisa Larkin and Nicole Herbst.

## FUNDING

This research was supported by NIMH R01-MH125833 and the Baszucki Brain Research Foundation to HP; Joe W. and Dorothy Dorsett Brown Foundation Neuroscience Initiative: AWD-001292, NIGMS P20-GM144041 to BG; NIAAA R24-AA019431 to KAG and R01-AA029023 to DMP.

## AUTHOR CONTRIBUTIONS

T.P. L.P, S.O., and O.A. contributed to data collection, T.P., A.Z., B.G., and H.P. analyzed the data, T.P., B.G, H.P., S.O., D.M.P., K.A.G. and K.F. contributed to data interpretation, T.P., B.G. and H.P. wrote the manuscript, D.M.P., K.A.G., A.Z., and K.F. contributed to manuscript preparation, T.P., B.G. and H.P. designed the studies.

## DATA AVAILABILITY

The data generated or analyzed during this study are available from the corresponding author upon reasonable request.

## STATEMENT ON CONFLICT OF INTEREST

The author(s) declare no competing interests.

## Notes

### Competing Interest Statement

The authors have declared no competing interest.

