## Supplemental Materials for "Cross-Species Evidence for Hippocampal CACNA1C as a Therapeutic Target for Alcohol Use Disorder"

**Tables S1-S4: Basic demographic information for all diagnosis groups**

Table S1: Basic demographic information for subjects with alcohol use disorder

| Age | Sex | Race | pH | PMI (hrs) | Sleep quality | ZT time | A-P Coordinates |
|---|---|---|---|---|---|---|---|
| | | | **Alcohol Use Disorder** | | | | |
| 40 | M | Black | 6.5 | 7 | *NA* | -2.42 | 31.9 |
| 32 | M | Black | 6.82 | 14 | *NA* | -6 | 26.5 |
| 29 | M | White | 6.75 | 27 | *NA* | 3.92 | 19.9 |
| 30 | M | White | 6.49 | 12 | *NA* | -3.7 | 33.1 |
| 38 | M | White | 6.68 | 25 | Decreased sleep | 7.02 | 27.8 |
| 38 | M | White | 6.42 | 6 | *NA* | *NA* | 26.5 |
| 49 | M | White | 6.88 | 24 | *NA* | *NA* | 25.2 |
| 34 | M | Black | 6.5 | 16 | *NA* | 17.77 | 27.8 |
| 36 | M | Black | 6.97 | 38.5 | Decreased sleep | -4.37 | 19.9 |
| 30 | M | Black | 6.8 | 21 | *NA* | *NA* | 23.9 |
| 42 | M | White | 6.86 | 30 | *NA* | 1.23 | 26.5 |
| 38 | M | White | 6.62 | 12 | *NA* | 17.03 | 39.5 |
| 31 | F | White | 6.18 | 12 | Decreased sleep | *NA* | 25.2 |
| 50 | M | White | 6.52 | 24 | *NA* | *NA* | 18.6 |
| 35 | F | Black | 6.43 | 26 | *NA* | *NA* | 16.0 |
| 22 | F | White | 6.12 | 12 | *NA* | *NA* | 21.2 |
| 56 | M | Black | 6.51 | 17 | *NA* | 13.14 | 33.1 |
| 28 | M | White | 6.3 | 26 | Decreased sleep | 3.67 | 21.2 |
| 52 | M | White | 6.81 | 8 | *NA* | 1.53 | 31.9 |
| 24 | M | White | 6.28 | 14 | *NA* | *NA* | 13.3 |
| mean ± SD 34.8±9.0 | 3F, 17M | 7B, 13W | 6.57±0.23 | 18.58±8.5 | | | 25.5±6.5 |

Table S2: Basic demographic information for subjects with major depressive disorder

| Age | Sex | Race | pH | PMI (hrs) | Sleep quality | ZT time | A-P Coordinates |
|---|---|---|---|---|---|---|---|
| | | | **Major Depressive Disorder** | | | | |
| 43 | M | White | 6.73 | 21 | Decreased sleep | 6.25 | 18.6 |
| 63 | F | White | 6.3 | 18 | Decreased sleep | 10.53 | 27.8 |
| 42 | M | White | 6.64 | 20 | *NA* | *NA* | 22.6 |
| 50 | F | White | 6.83 | 23 | Decreased sleep | 6 | 27.8 |
| 34 | F | White | 6.27 | 24 | Decreased sleep | NA | 29.2 |
| 55 | M | White | 6.64 | 29 | Decreased sleep | 2.08 | 27.8 |
| 33 | M | White | 6.79 | 18 | Increased sleep | *NA* | 26.5 |
| 53 | M | White | 6.73 | 29 | Decreased sleep | 4.25 | 27.8 |
| 44 | F | White | 6.71 | 29 | *NA* | 2.83 | 23.9 |
| 59 | M | White | 6.22 | 27 | Decreased sleep | 8.47 | 23.9 |
| 46 | M | Black | 6.26 | 17 | Decreased sleep | 13.47 | 33.1 |
| 36 | F | White | 6.84 | 25 | Increased sleep | 7.12 | 27.8 |
| 35 | M | White | 6.96 | 11 | Decreased sleep | 11.33 | 21.2 |
| 41 | M | White | 6.24 | 19 | Increased sleep | *NA* | 23.9 |
| 56 | F | White | 5.61 | 38 | Decreased sleep | *NA* | 23.9 |
| 62 | M | Black | 6.06 | 22 | Increased sleep | *NA* | 17.2 |
| 20 | M | White | 6.73 | 20 | Decreased sleep | *NA* | 34.6 |
| 59 | M | Black | 6.6 | 31 | Decreased sleep | *NA* | 23.9 |
| 48 | M | White | 6.41 | 27 | Normal sleep | 2.78 | 31.9 |
| 61 | M | White | 6.74 | 25 | Decreased sleep | 6.33 | 26.6 |
| mean ± SD 47±11.4 | 6F, 14M | 3B, 17W | 6.5±0.32 | 23.7±5.9 | | | 26.0±4.4 |

Table S3: Basic demographic information for subjects with comorbid alcohol use disorder and major depressive disorder

| Age | Sex | Race | pH | PMI (hrs) | Sleep quality | ZT time | A-P Coordinates |
|---|---|---|---|---|---|---|---|
| | | | **Comorbid AUD and MDD** | | | | |
| 36 | M | White | 6.72 | 15 | Decreased sleep | 15.33 | 19.9 |
| 37 | M | White | 6.89 | 19 | *NA* | 11.08 | 31.9 |
| 41 | F | White | 6.55 | 17 | Decreased sleep | *NA* | 25.2 |
| 47 | F | White | 6.65 | 9 | *NA* | -3.5 | 27.8 |
| 48 | F | White | 6.13 | 24 | Increased sleep | 2.25 | 21.2 |
| 44 | M | White | 6.77 | 20 | Decreased sleep | 8.16 | 39.5 |
| 29 | F | White | 6.47 | 29 | Decreased sleep | 4.58 | 26.5 |
| 59 | F | White | 6.8 | 24 | *NA* | 6.68 | 34.6 |
| 40 | M | White | 6.66 | 26 | *NA* | 2.16 | 26.5 |
| 45 | M | White | 6.29 | 24 | *NA* | *NA* | 31.9 |
| 20 | M | Black | 6.21 | 10 | Increased sleep | *NA* | 22.6 |
| 35 | M | White | 6.81 | 24 | *NA* | 5.62 | 29.2 |
| 34 | M | White | 6.33 | 17 | *NA* | 13.25 | 14.6 |
| 37 | M | White | 6.59 | 18 | *NA* | 14 | 21.2 |
| 63 | F | White | 6.32 | 24 | Decreased sleep | 5.63 | 22.6 |
| 62 | M | Black | 6.52 | 17 | Decreased sleep | 15.98 | 23.9 |
| 54 | M | White | 6.54 | 38 | Decreased sleep | 8.5 | 17.2 |
| 48 | F | Black | 5.87 | 17 | Decreased sleep | *NA* | 21.2 |
| 43 | M | White | 6.6 | 20 | Decreased sleep | 13.65 | 21.2 |
| 42 | F | White | 6.84 | 12 | Decreased sleep | -4.75 | 17.5 |
| 40 | M | White | 6.62 | 21 | Normal sleep | 8.5 | 25.2 |
| 58 | M | Black | 6.11 | 37 | *NA* | *NA* | 27.8 |
| 30 | M | White | 6.91 | 18 | *NA* | 12.42 | 34.6 |
| 62 | M | White | 6.7 | 5 | *NA* | 0.37 | 39.5 |
| mean ± SD 43.9±11.1 | 8F, 16M | 4B, 20W | 6.54±0.27 | 20.2±7.6 | | | 26.0±6.7 |

Table S4: Basic demographic information for unaffected control subjects

| Age | Sex | Race | pH | PMI (hrs) | Sleep quality | ZT time | A-P Coordinates |
|---|---|---|---|---|---|---|---|
| | | | **Unaffected Control Subjects** | | | | |
| 62 | F | White | 6.34 | 27.5 | Normal sleep | 0.67 | 21.2 |
| 54 | M | Black | 6.53 | 19 | Normal sleep | 9.67 | 29.2 |
| 52 | M | White | 6.28 | 17 | Normal sleep | 13.15 | 23.9 |
| 30 | M | Black | 6.98 | 19 | Normal sleep | 13.92 | 27.8 |
| 48 | M | Black | 6.98 | 9 | Normal sleep | 0.28 | 18.6 |
| 51 | F | Black | 6.3 | 22 | Normal sleep | 8.25 | 23.9 |
| 49 | F | Black | 6.57 | 29 | Normal sleep | 5 | 22.6 |
| 38 | F | White | 5.93 | 13 | Normal sleep | -1.88 | 23.9 |
| 44 | F | Black | 6.72 | 32 | Decreased sleep | 2.17 | 23.9 |
| 44 | M | White | 6.6 | 24.32 | Normal sleep | *NA* | 39.5 |
| 28 | M | Black | 6.32 | 35.3 | Normal sleep | -2.22 | 17.2 |
| 42 | F | White | 6.6 | 21 | Normal sleep | 10.72 | 23.9 |
| 31 | M | White | 6.78 | 14.15 | Normal sleep | 15 | 29.2 |
| 51 | M | White | 6.76 | 17 | Decreased sleep | 16.17 | 10.7 |
| 29 | F | Black | 6.64 | 25.5 | Normal sleep | *NA* | 39.5 |
| 35 | M | Black | 6.26 | 21 | Normal sleep | 9.5 | 21.2 |
| 49 | M | White | 6.71 | 9.75 | Decreased sleep | -1.67 | 27.8 |
| 17 | M | White | 6.66 | 22.75 | Normal sleep | -4.27 | 31.9 |
| 51 | M | White | 6.3 | | Normal sleep | *NA* | 30.5 |
| 59 | M | White | 6.47 | 23.75 | Normal sleep | 6.75 | 23.9 |
| mean ± SD 43.2±11.42 | 7F, 13M | 9B, 11W | 6.5±0.25 | 21.2±6.8 | | | 25.5±6.9 |

**Tables S5-S8: Substance use information for all diagnostic groups**

Table S5: Substance use information for subjects with alcohol use disorder

| case | Onset of AUD (age) | Duration of AUD (yrs) | Ethanol in toxicology report | Cocaine in toxicology report | Opioids in toxicology report | Opioid dependence | Smoker | Nicotine rating | Alcohol rating | Cannabis history |
|---|---|---|---|---|---|---|---|---|---|---|
| 40M | 14 | 26 | Yes | Yes | No | No | Yes | 2 | 4 | No |
| 32M | *NA* | *NA* | Yes | Yes | No | No | Yes | 3 | 2 | Yes |
| 29M | 14 | 15 | No | No | No | No | Yes | 4 | 4 | Yes |
| 30M | 20 | 10 | Yes | No | No | No | Yes | *NA* | 4 | Yes |
| 38M | 20 | 18 | No | No | Yes | Yes | No | 0 | 2 | Yes |
| 38M | 16 | 22 | Yes | No | Yes | Yes | Yes | 3 | 4 | Yes |
| 49M | 18 | 31 | Yes | No | No | Yes | Yes | 4 | 4 | Yes |
| 34M | 16 | 18 | No | Yes | No | No | Yes | 4 | 4 | Yes |
| 36M | 16 | 20 | Yes | No | No | No | Yes | 2 | 4 | Yes |
| 30M | 19 | 11 | No | Yes | No | No | Yes | 2 | 3 | Yes |
| 42M | 15 | 27 | No | No | No | No | Yes | 4 | 4 | No |
| 38M | 14 | 24 | Yes | Yes | Yes | Yes | Yes | 4 | 4 | Yes |
| 31F | 16 | 15 | No | No | No | No | Yes | 4 | 4 | Yes |
| 50M | 14 | 36 | Yes | Yes | No | Yes | No | 0 | 4 | No |
| 35F | 15 | 20 | No | Yes | No | No | No | 0 | 4 | Yes |
| 22F | 12 | 10 | Yes | No | No | No | Yes | 4 | 4 | Yes |
| 56M | 26 | 30 | No | Yes | No | No | No | 0 | 4 | Yes |
| 28M | *NA* | *NA* | No | No | Yes | Yes | Yes | 2 | 4 | Yes |
| 52M | 16 | 35 | Yes | No | No | No | Yes | 4 | 4 | Yes |
| 24M | 21 | 3 | Yes | Yes | Yes | Yes | Yes | 4 | 4 | Yes |
| mean ± SD 34.8±9.0/ 3F, 17M | | | | | | | | | | |

Table S6: Substance use information for subjects with major depressive disorder

| case | Onset of AUD (age) | Duration of AUD (yrs) | Ethanol in toxicology report | Cocaine in toxicology report | Opioids in toxicology report | Opioid dependence | Smoker | Nicotine rating | Alcohol rating | Cannabis history |
|---|---|---|---|---|---|---|---|---|---|---|
| 43M | None | 0 | No | No | No | No | No | 0 | 0 | No |
| 63F | None | 0 | No | No | No | No | Yes | 4 | 1 | No |
| 42M | None | 0 | No | No | No | No | No | 0 | 0 | No |
| 50F | None | 0 | No | No | No | No | Yes | 4 | 1 | No |
| 34F | None | 0 | No | No | No | No | No | 0 | 0 | No |
| 55M | None | 0 | No | No | No | No | Yes | 3 | 0 | No |
| 33M | None | 0 | No | No | No | No | No | 0 | 1 | No |
| 53M | None | 0 | No | No | No | No | No | 0 | 1 | No |
| 44F | None | 0 | No | No | No | No | Yes | 3 | 0 | No |
| 59M | None | 0 | No | No | No | No | Yes | 4 | 2 | No |
| 46M | None | 0 | No | No | No | No | No | 0 | 2 | No |
| 36F | None | 0 | No | No | No | No | Yes | 4 | 2 | No |
| 35M | None | 0 | No | No | No | No | No | 0 | 1 | No |
| 41M | None | 0 | No | No | No | No | Yes | 4 | 1 | No |
| 56F | None | 0 | No | No | No | No | Yes | 4 | 2 | No |
| 62M | None | 0 | No | No | No | No | Yes | 3 | 1 | No |
| 20M | None | 0 | No | No | No | No | No | 0 | 1 | No |
| 59M | None | 0 | No | No | No | No | No | 0 | 1 | No |
| 48M | None | 0 | No | No | No | No | Yes | 3 | 1 | No |
| 61M | None | 0 | No | No | Yes | No | Yes | 2 | 1 | No |
| mean ± SD 47±11.4/ 6F, 14M | | | | | | | | | | |

Table S7: Substance use information for subjects with comorbid alcohol use disorder and major depressive disorder

| case | Onset of AUD (age) | Duration of AUD (yrs) | Ethanol in toxicology report | Cocaine in toxicology report | Opioids in toxicology report | Opioid dependence | Smoker | Nicotine rating | Alcohol rating | Cannabis history |
|---|---|---|---|---|---|---|---|---|---|---|
| 36M | 8 | 28 | Yes | Yes | No | No | Yes | 4 | 4 | Yes |
| 37M | 14 | 23 | Yes | No | No | No | Yes | 4 | 4 | Yes |
| 41F | *NA* | *NA* | Yes | No | No | Yes | No | *NA* | 1 | No |
| 47F | 27 | 20 | No | No | No | Yes | Yes | 4 | 4 | Yes |
| 48F | 45 | 3 | Yes | No | No | No | Yes | 4 | 4 | No |
| 44M | 34 | 10 | No | No | No | No | Yes | 4 | 4 | No |
| 29F | *NA* | *NA* | No | No | Yes | Yes | Yes | 4 | 4 | No |
| 59F | 34 | 25 | No | No | Yes | No | Yes | 4 | 4 | No |
| 40M | 13 | 27 | Yes | Yes | No | No | Yes | *NA* | 4 | Yes |
| 45M | *NA* | *NA* | Yes | No | No | No | Yes | 4 | 4 | No |
| 20M | 19 | 1 | No | No | No | No | No | 0 | 3 | Yes |
| 35M | *NA* | *NA* | No | No | No | No | No | 0 | 4 | No |
| 34M | 14 | 20 | Yes | No | No | No | Yes | 2 | 3 | Yes |
| 37M | 14 | 23 | Yes | No | Yes | Yes | Yes | 3 | 4 | No |
| 63F | 38 | 25 | Yes | No | No | Yes | Yes | 4 | 4 | No |
| 62M | 42 | 20 | Yes | No | No | No | Yes | 2 | 4 | No |
| 54M | 24 | 30 | No | No | No | No | Yes | 4 | 4 | No |
| 48F | *NA* | *NA* | Yes | No | Yes | Yes | Yes | 3 | 4 | Yes |
| 43M | *NA* | *NA* | Yes | No | No | Yes | Yes | 4 | 1 | No |
| 42F | *NA* | *NA* | No | No | No | Yes | Yes | 1 | 3 | Yes |
| 40M | *NA* | *NA* | No | No | No | No | Yes | 4 | 2 | Yes |
| 58M | 23 | 35 | Yes | No | Yes | Yes | Yes | 4 | 4 | No |
| 30M | *NA* | *NA* | Yes | No | No | No | Yes | 3 | 3 | Yes |
| 62M | *NA* | 20 | No | No | No | No | Yes | 4 | 4 | No |
| mean ± SD 43.9±11.1/ 8F, 16M | | | | | | | | | | |

Table S8: Substance use information for unaffected control subjects

| case | Onset of AUD (age) | Duration of AUD (yrs) | Ethanol in toxicology report | Cocaine in toxicology report | Opioids in toxicology report | Opioid dependence | Smoker | Nicotine rating | Alcohol rating | Cannabis history |
|---|---|---|---|---|---|---|---|---|---|---|
| 62F | None | 0 | No | No | No | No | No | 0 | 1 | No |
| 54M | None | 0 | No | No | No | No | No | 4 | 3 | No |
| 52M | None | 0 | No | No | No | No | No | 0 | 0 | No |
| 30M | None | 0 | No | No | No | No | No | 1 | 1 | No |
| 48M | None | 0 | No | No | No | No | No | 4 | 1 | No |
| 51F | None | 0 | No | No | No | No | Yes | 4 | 1 | Yes |
| 49F | None | 0 | No | No | No | No | No | 0 | 3 | No |
| 38F | None | 0 | No | No | No | No | No | 0 | 0 | No |
| 44F | None | 0 | No | No | No | No | Yes | 2 | 1 | No |
| 44M | None | 0 | No | No | No | No | Yes | 4 | 1 | No |
| 28M | None | 0 | No | No | No | No | No | 0 | 1 | No |
| 42F | None | 0 | No | No | No | No | Yes | 4 | 2 | Yes |
| 31M | None | 0 | No | No | No | No | Yes | 4 | 2 | No |
| 51M | None | 0 | No | No | No | No | Yes | 4 | 2 | No |
| 29F | None | 0 | No | No | No | No | Yes | 2 | 2 | Yes |
| 35M | None | 0 | No | No | No | No | Yes | 2 | 2 | No |
| 49M | None | 0 | No | No | No | No | Yes | 4 | 2 | Yes |
| 17M | None | 0 | Yes | No | No | No | No | 0 | 2 | No |
| 51M | None | 0 | No | No | No | No | No | 1 | 2 | Yes |
| 59M | None | 0 | No | No | No | No | Yes | 3 | 1 | No |
| mean ± SD 43.2±11.42/ 7F, 13M | | | | | | | | | | |

**Tables S9-S12: Mood disorder-related information for all diagnostic groups**

Table S9: Mood disorder-related information for subjects with alcohol use disorders

| case | Suicide | Depression severity | Duration of MDD (yrs.) | Antidepressants in blood at death | Antidepressants/ month (grams) | Mania | Psychosis | Antipsychotics |
|---|---|---|---|---|---|---|---|---|
| | | | | **Alcohol Use Disorder** | | | | |
| 62F | No | None | 0 | No | 0 | No | No | No |
| 54M | No | None | 0 | No | 0 | No | No | No |
| 52M | No | Mild | 0 | No | 0 | No | No | No |
| 30M | No | None | 0 | No | 0 | No | No | No |
| 48M | No | *NA* | 0 | Yes | 4.5 | No | No | No |
| 51F | No | None | 0 | No | 0 | No | No | No |
| 49F | Yes | None | 0 | No | 0 | No | No | No |
| 38F | No | None | 0 | No | 0 | No | No | No |
| 44F | No | None | 0 | No | 0 | No | No | No |
| 44M | No | None | 0 | No | 0 | No | No | No |
| 28M | No | None | 0 | No | 0 | No | No | No |
| 42F | No | None | 0 | No | 0 | No | No | No |
| 31M | No | *NA* | *NA* | Yes | 10.2 | No | Yes | Yes |
| 51M | Yes | *NA* | *NA* | Yes | 4.8 | No | No | No |
| 29F | No | None | *NA* | No | 0 | No | No | No |
| 35M | No | None | 0 | No | 0 | No | No | No |
| 49M | No | None | 0 | No | 0 | No | No | No |
| 17M | No | None | 0 | No | 0 | No | No | No |
| 51M | No | *NA* | *NA* | No | 0 | No | No | No |
| 59M | Yes | *NA* | *NA* | Yes | 2.6 | No | No | No |
| mean ± SD 34.8±9.0/ 3F, 17M | | | | | | | | |

Table S10: Mood disorder-related information for subjects with major depressive disorder

| case | Suicide | Depression severity | Duration of MDD (yrs.) | Antidepressants in blood at death | Antidepressants/ month (grams) | Mania | Psychosis | Antipsychotics |
|---|---|---|---|---|---|---|---|---|
| | | | | **Major Depressive Disorder** | | | | |
| 43M | Yes | Moderate | 15 | No | 0 | No | Yes | No |
| 63F | No | Moderate | *NA* | Yes | 3 | No | No | No |
| 42M | Yes | Severe | *NA* | Yes | 0.8 | No | No | No |
| 50F | Yes | Moderate | *NA* | Yes | 0 | No | Yes | No |
| 34F | Yes | Severe | *NA* | Yes | 3 | No | No | Yes |
| 55M | Yes | *NA* | *NA* | Yes | 3.2 | No | No | No |
| 33M | Yes | Moderate | *NA* | Yes | 16.7 | No | No | Yes |
| 53M | Yes | *NA* | *NA* | Yes | 4 | No | No | No |
| 44F | Yes | *NA* | 12 | Yes | 8.7 | No | No | No |
| 59M | Yes | *NA* | 0.08 | Yes | 3 | No | No | No |
| 46M | No | Mild | 1 | No | 0 | No | No | No |
| 36F | No | Mild | *NA* | No | 0 | No | No | No |
| 35M | No | Moderate | 1.5 | No | 0 | No | No | No |
| 41M | No | Moderate | 7 | No | 0 | No | No | No |
| 56F | No | *NA* | 1 | No | 0 | No | No | No |
| 62M | No | Mild | 49 | No | 0 | No | No | No |
| 20M | Yes | Moderate | 1.33 | No | 0 | No | No | No |
| 59M | Yes | *NA* | *NA* | Yes | 0.4 | No | No | No |
| 48M | No | *NA* | *NA* | Yes | 2.7 | No | No | No |
| 61M | Yes | Severe | 47 | Yes | 3.6 | No | No | No |
| mean ± SD 47±11.4/ 6F, 14M | | | | | | | | |

Table S11: Mood disorder-related information for subjects with comorbid alcohol use disorder and major depressive disorder

| case | Suicide | Depression severity | Duration of MDD (yrs.) | Antidepressants in blood at death | Antidepressants/ month (grams) | Mania | Psychosis | Antipsychotics |
|---|---|---|---|---|---|---|---|---|
| **Comorbid AUD and MDD** | | | | | | | | |
| 36M | Yes | Severe | 16 | Yes | 3 | No | Yes | No |
| 37M | Yes | Moderate | 18 | Yes | 2 | No | No | No |
| 41F | Yes | Moderate | 14 | Yes | 3 | No | No | No |
| 47F | No | Moderate | 20 | Yes | 0 | No | No | No |
| 48F | Yes | Severe | 9 | Yes | 4.3 | No | No | No |
| 44M | Yes | Mild | 8 | Yes | 12 | No | No | No |
| 29F | No | Severe | 18 | Yes | 4 | No | Yes | No |
| 59F | Yes | Moderate | 25 | Yes | 3 | No | No | No |
| 40M | Yes | *NA* | 24 | No | 0 | No | No | No |
| 45M | Yes | Moderate | 25 | Yes | 2.6 | No | No | No |
| 20M | Yes | *NA* | 1 | Yes | 4.5 | No | No | No |
| 35M | Yes | *NA* | 18 | Yes | 8.8 | No | No | Yes |
| 34M | Yes | Mild | 8 | No | 0 | No | No | No |
| 37M | Yes | Severe | 27 | Yes | 2.3 | No | Yes | Yes |
| 63F | No | Moderate | 30 | Yes | 1.5 | No | Yes | Yes |
| 62M | No | Severe | 30 | No | 0 | No | No | No |
| 54M | No | *NA* | 40 | Yes | 3 | No | No | No |
| 48F | No | Moderate | 19 | Yes | 4.2 | No | No | No |
| 43M | Yes | Moderate | 24 | Yes | 0.8 | No | No | No |
| 42F | No | Moderate | *NA* | Yes | 6.4 | No | Yes | Yes |
| 40M | No | Severe | 30 | Yes | 3 | No | Yes | No |
| 58M | No | *NA* | 20 | Yes | 4.5 | No | No | No |
| 30M | Yes | Moderate | *NA* | No | 0 | No | No | No |
| 62M | Yes | Severe | 25 | Yes | 5.7 | No | No | No |
| mean ± SD 43.9±11.1/ 8F, 16M | | | | | | | | |

Table S12: Mood disorder-related information for unaffected control subjects

| case | Suicide | Depression severity | Duration of MDD (yrs.) | Antidepressants in blood at death | Antidepressants/ month (grams) | Mania | Psychosis | Antipsychotics |
|---|---|---|---|---|---|---|---|---|
| **Unaffected Control Subjects** | | | | | | | | |
| 62F | No | None | 0 | No | 0 | No | No | No |
| 54M | No | None | 0 | No | 0 | No | No | No |
| 52M | No | None | 0 | No | 0 | No | No | No |
| 30M | No | None | 0 | No | 0 | No | No | No |
| 48M | No | None | 0 | No | 0 | No | No | No |
| 51F | No | None | 0 | No | 0 | No | No | No |
| 49F | No | None | 0 | No | 0 | No | No | No |
| 38F | No | None | 0 | No | 0 | No | No | No |
| 44F | No | Mild | 0 | No | 0 | No | No | No |
| 44M | No | None | 0 | No | 0 | No | No | No |
| 28M | No | None | 0 | No | 0 | No | No | No |
| 42F | No | Mild | 0 | No | 0 | No | No | No |
| 31M | No | None | 0 | No | 0 | No | No | No |
| 51M | No | None | 0 | No | 0 | No | No | No |
| 29F | No | Mild | 0 | No | 0 | No | No | No |
| 35M | No | None | 0 | No | 0 | No | No | No |
| 49M | No | None | 0 | No | 0 | No | No | No |
| 17M | No | None | 0 | No | 0 | No | No | No |
| 51M | No | None | 0 | No | 0 | No | No | No |
| 59M | No | None | 0 | No | 0 | No | No | No |
| mean ± SD 43.2±11.42/ 7F, 13M | | | | | | | | |

**Tables S13-S16: Other relevant demographic information for all diagnostic groups**

Table S13: Other relevant information for subjects with alcohol use disorders

| case | Anxiety disorders | Personality disorders | Obsessive compulsive disorder | Lithium | Lithium/ mo (grams) | Calcium channel blockers | Antipsychotics last month of life (CPZ eq./grams) | Valproic acid last month of life (grams) |
|---|---|---|---|---|---|---|---|---|
| | | | | **Alcohol Use Disorder** | | | | |
| 62F | None | Antisocial | No | None | 0 | No | 0 | 0 |
| 54M | None | Antisocial | No | None | 0 | No | 0 | 0 |
| 52M | None | Antisocial | No | None | 0 | No | 0 | 0 |
| 30M | None | None | No | None | 0 | No | 0 | 0 |
| 48M | None | None | No | None | 0 | No | 0 | 0 |
| 51F | None | Antisocial | No | None | 0 | No | 0 | 0 |
| 49F | None | Antisocial | No | None | 0 | No | 0 | 0 |
| 38F | None | Antisocial | No | None | 0 | No | 0 | 0 |
| 44F | None | Antisocial | No | None | 0 | No | 0 | 0 |
| 44M | None | None | No | None | 0 | No | 0 | 0 |
| 28M | None | None | No | None | 0 | No | 0 | 0 |
| 42F | PTSD | Antisocial | No | None | 0 | No | 0 | 0 |
| 31M | GAD | Borderline/ Antisocial | No | None | 0 | No | 6 | 0 |
| 51M | | Antisocial | No | None | 0 | No | 0 | 0 |
| 29F | None | None | No | None | 0 | No | 0 | 0 |
| 35M | None | None | No | None | 0 | No | 0 | 0 |
| 49M | None | None | No | None | 0 | No | 0 | 0 |
| 17M | None | None | No | None | 0 | No | 0 | 0 |
| 51M | None | Antisocial | No | None | 0 | No | 0 | 0 |
| 59M | None | Borderline | No | None | 0 | No | 0 | 0 |
| mean ± SD 34.8±9.0/ 3F, 17M | | | | | | | | |

Table S14: Other relevant information for subjects with major depressive disorder

| case | Anxiety disorders | Personality disorders | Obsessive compulsive disorder | Lithium | Lithium/ mo (grams) | Calcium channel blockers | Antipsychotics last month of life (CPZ eq./grams) | Valproic acid last month of life (grams) |
|---|---|---|---|---|---|---|---|---|
| **Major Depressive Disorder** | | | | | | | | |
| 43M | None | None | Yes | None | 0 | No | 0 | 0 |
| 63F | None | Schizoid | No | None | 0 | No | 0 | 0 |
| 42M | None | None | No | None | 0 | No | 0 | 0 |
| 50F | None | None | No | None | 0 | No | 0 | 0 |
| 34F | Panic | None | No | None | 0 | No | 1.1 | 7.5 |
| 55M | None | None | No | None | 0 | No | 0 | 0 |
| 33M | None | Dependent | No | Yes | 12 | No | 6.8 | 0 |
| 53M | None | None | No | None | 0 | *NA* | 0 | 0 |
| 44F | None | None | No | None | 0 | No | 0 | 0 |
| 59M | GAD | None | No | None | 0 | No | 0 | 0 |
| 46M | None | Schizoid | No | None | 0 | No | 0 | 0 |
| 36F | Panic | None | No | None | 0 | Yes | 0 | 0 |
| 35M | None | None | Yes | None | 0 | No | 0 | 0 |
| 41M | None | None | No | None | 0 | No | 0 | 0 |
| 56F | GAD | None | No | None | 0 | No | 0 | 0 |
| 62M | None | None | No | None | 0 | Yes | 0 | 0 |
| 20M | Adjustment | None | No | None | 0 | No | 0 | 0 |
| 59M | None | None | No | None | 0 | No | 0 | 0 |
| 48M | *NA* | *NA* | *NA* | None | 0 | No | 0 | 0 |
| 61M | None | None | No | None | 0 | No | 0 | 0 |
| mean ± SD 47±11.4/ 6F, 14M | | | | | | | | |

Table S15: Other relevant information for subjects with comorbid alcohol use disorder and major depressive disorder

| case | Anxiety disorders | Personality disorders | Obsessive compulsive disorder | Lithium | Lithium/ mo (grams) | Calcium channel blockers | Antipsychotics last month of life (CPZ eq.) | Valproic acid last month of life (grams) |
|---|---|---|---|---|---|---|---|---|
| **Comorbid AUD and MDD** | | | | | | | | |
| 36M | None | None | No | None | 0 | No | 0 | 0 |
| 37M | None | Dependent | No | None | 0 | No | 0 | 0 |
| 41F | None | None | No | None | 0 | No | 0 | 0 |
| 47F | Phobia | Histrionic | No | None | 0 | No | 0 | 0 |
| 48F | None | None | No | None | 0 | No | 0 | 0 |
| 44M | PTSD | None | No | None | 0 | No | 0 | 0 |
| 29F | None | Borderline | No | None | 0 | No | 0 | 0 |
| 59F | Panic | None | No | None | 0 | No | 0 | 0 |
| 40M | None | None | No | None | 0 | No | 0 | 0 |
| 45M | None | Borderline | No | None | 0 | No | 0 | 0 |
| 20M | GAD | None | No | None | 0 | No | 0 | 0 |
| 35M | None | Mixed | No | None | 0 | No | 2 | 0 |
| 34M | None | Borderline | No | None | 0 | No | 0 | 0 |
| 37M | None | Borderline / Antisocial | No | None | 0 | No | 1.8 | 0 |
| 63F | None | Dependent | No | None | 0 | No | 0.9 | 0 |
| 62M | None | Borderline | No | None | 0 | No | 0 | 0 |
| 54M | None | None | No | None | 0 | No | 0 | 0 |
| 48F | None | None | No | None | 0 | No | 0 | 0 |
| 43M | None | None | No | None | 0 | No | 0 | 0 |
| 42F | None | None | No | None | 0 | No | 3.6 | 0 |
| 40M | None | Dependent | No | None | 0 | No | 0 | 0 |
| 58M | None | None | No | None | 0 | No | 0 | 0 |
| 30M | None | Borderline | No | None | 0 | No | 0 | 0 |
| 62M | *NA* | *NA* | *NA* | None | 0 | No | 0 | 0 |
| mean ± SD 43.9±11.1/ 8F, 16M | | | | | | | | |

Table S16: Other relevant information for unaffected control subjects

| case | Anxiety disorders | Personality disorders | Obsessive compulsive disorder | Lithium | Lithium/ mo (grams) | Calcium channel blockers | Antipsychotics last month of life (CPZ eq.) | Valproic acid last month of life (grams) |
|---|---|---|---|---|---|---|---|---|
| **Unaffected Control Subjects** | | | | | | | | |
| 62F | *NA* | *NA* | *NA* | None | 0 | No | 0 | 0 |
| 54M | None | None | Yes | None | 0 | No | 0 | 0 |
| 52M | *NA* | *NA* | *NA* | None | 0 | No | 0 | 0 |
| 30M | None | None | No | None | 0 | No | 0 | 0 |
| 48M | None | None | No | None | 0 | No | 0 | 0 |
| 51F | None | None | Yes | None | 0 | No | 0 | 0 |
| 49F | *NA* | *NA* | *NA* | None | 0 | No | 0 | 0 |
| 38F | *NA* | *NA* | *NA* | None | 0 | No | 0 | 0 |
| 44F | None | None | Yes | None | 0 | *NA* | 0 | 0 |
| 44M | None | None | Yes | None | 0 | No | 0 | 0 |
| 28M | None | None | No | None | 0 | No | 0 | 0 |
| 42F | None | None | Yes | None | 0 | No | 0 | 0 |
| 31M | *NA* | *NA* | *NA* | None | 0 | No | 0 | 0 |
| 51M | *NA* | *NA* | *NA* | None | 0 | No | 0 | 0 |
| 29F | None | None | No | None | 0 | No | 0 | 0 |
| 35M | None | None | No | None | 0 | Yes | 0 | 0 |
| 49M | None | None | No | None | 0 | No | 0 | 0 |
| 17M | *NA* | *NA* | *NA* | None | 0 | No | 0 | 0 |
| 51M | None | Paranoid | No | None | 0 | No | 0 | 0 |
| 59M | None | None | Yes | None | 0 | No | 0 | 0 |
| mean ± SD 43.2±11.42/ 7F, 13M | | | | | | | | |

***NA* = information not available**
**GAD = generalized anxiety disorder**
**PTSD = post-traumatic stress disorder**

**Table S17. QPCR Primers**

| Probes | Forward | Reverse |
|---|---|---|
| **EXPERIMENTAL** | | |
| *CACNA1C* | TGATTCCAACGCCACCAATTC | GAGGAGTCCATAGGCGATTACT |
| **LOADING CONTROLS** | | |
| *Actb* | GTCATTCCAAATATGAGATGCGT | GCTATCACCTCCCCTGTGTG |
| *Ppia* | ATGGTCAACCCCACCGTGTTCTTCG | CGTGTGAAGTCACCACCCTGACACA |
| *Gapdh* | TCGACAGTCAGCCGCATCT | AGTTAAAAGCAGCCCTGGTGA |
| *B2m* | GTGGGATCGAGACATGTAAGC | AGCAAGCAAGCAGAATTTGGAAT |
